# Transcranial magnetic stimulation of right dorsolateral prefrontal cortex does not affect associative retrieval in healthy young or older adults

**DOI:** 10.1101/2021.01.28.428662

**Authors:** Paul F. Hill, Erin D. Horne, Joshua D. Koen, Michael D. Rugg

## Abstract

We examined whether post-retrieval monitoring processes supporting memory performance are more resource limited in older adults relative to younger individuals. We predicted that older adults would be more susceptible to an experimental manipulation that reduced the neurocognitive resources available to support post-retrieval monitoring. Young and older adults received transcranial magnetic stimulation (TMS) to the right dorsolateral prefrontal cortex (DLPFC) or a vertex control site during an associative recognition task. The right DLPFC was selected as a TMS target because the region is held to be a key member of a network of regions engaged during retrieval monitoring and is readily accessible to administration of TMS. We predicted that TMS to the right DLPFC would lead to reduced associative recognition accuracy, and that this effect would be more prominent in older adults. The results did not support this prediction. Recognition accuracy was significantly reduced in older adults relative to their younger counterparts but the magnitude of this age difference was unaffected following TMS to the right DLPFC or vertex. These findings suggest that TMS to the right DLPFC was insufficient to deplete the neurocognitive resources necessary to support post-retrieval monitoring.

---

Episodic memory, or the ability to recollect specific details about a unique event, declines with advancing age (Koen & Yonelinas, 2014; Nilsson, 2003; Old & Naveh-Benjamin, 2008; Park et al., 1996; Schoemaker et al., 2014). Successful recollection is assumed to rely on multiple, interacting cognitive processes (Eichenbaum et al., 2007; Yonelinas, 2002; Rugg & Wilding, 2000 for review). Here, we focus on processes held to be engaged during evaluation of the outcome of a retrieval attempt (*post-retrieval monitoring*). Specifically, we examined whether older adults are more vulnerable to an experimental manipulation that was hypothesized to reduce the resources available for post-retrieval monitoring. To address this question, we used transcranial magnetic stimulation (TMS) to temporarily deplete cognitive resources dependent on the right dorsolateral prefrontal cortex (DLPFC) – a region held to be a key member of the brain network supporting monitoring – during an associative retrieval task.

Post-retrieval monitoring refers to a class of control processes engaged to evaluate the outcome of a retrieval attempt in relation to a behavioral goal (Burgess & Shallice, 1996; Rugg, 2004). For example, when asked to remember one’s most recent trip to a favorite restaurant, several competing memories are likely brought to mind. Accurate retrieval requires the selection and maintenance of the appropriate retrieval goal (memory for the most recent visit), and then the monitoring and evaluation of retrieved content in light of this goal (discriminating between memories for the most recent and prior visits; Rugg & Wilding, 2000). Importantly, post-retrieval monitoring is widely assumed to be supported by domain-general executive control processes which are shared with other cognitive operations (Cocci et al., 2013; Duncan, 2010; Hayama et al., 2008; Hayama & Rugg, 2009). We therefore predicted that behavioral outcomes associated with successful monitoring would be impaired by experimental manipulations aimed at interfering with these domain-general processes.

Neuropsychological studies of patients with lesions of the prefrontal cortex (PFC) were among the first to provide a neurobiological basis for post-retrieval monitoring. Compared to healthy control subjects, patients with frontal lesions were reported to display elevated false alarm rates on tests of recognition memory, a finding held to be diagnostic of monitoring failure (Burgess & Shallice; Johnson et al., 1997; Schacter et al., 1996). Concordant with these early findings, more recent studies using functional magnetic resonance imaging (fMRI) have consistently reported retrieval ‘monitoring effects’ – operationalized as enhanced brain activity elicited by weak relative to strong memory signals – in right and, to a lesser extent, left DLPFC as well as in the anterior cingulate cortex (e.g. Achim & Lepage, 2005; de Chastelaine et al., 2016; Henson et al., 1999, 2000; Horne et al., 2020; Wang et al., 2015).

The executive control processes assumed to support post-retrieval monitoring are generally considered highly vulnerable to advancing age (e.g., Braver & Barch, 2002; Buckner, 2004; Grady, 2012; Hedden & Gabrieli, 2004). However, only a handful of studies have contrasted fMRI retrieval monitoring effects between healthy young and older adults. The findings from these studies are mixed, with some studies reporting a null effect of age (de Chastelaine et al., 2016; Dulas & Duarte, 2014; Giovanello et al., 2010; Wang et al., 2015), and others reporting attenuated monitoring effects in older relative to young adults (Duarte et al., 2010; McDonough et al., 2013; Mitchell et al., 2013; see Horne et al., 2020 for detailed discussion of these discrepant findings). Of importance, correlations between frontal monitoring effects and memory performance have been reported to be both highly robust and age invariant (de Chastelaine et al., 2016; Horne et al., 2020; Wang et al., 2015), suggesting that age does not moderate the functional relationship between the effects and memory accuracy.

Recently, we (Horne et al., 2020) proposed that a key factor determining whether age differences are observed in the capacity to effectively engage retrieval monitoring is the availability of the neurocognitive resources required to support monitoring operations. According to the CRUNCH model (Reuter-Lorenz & Cappell, 2008), for example, recruitment of task-related neural resources will track task demands until a resource limit is reached, at which point (the ‘crunch point’) task performance will suffer. According to this model, other things equal, older adults will reach this resource limit at lower levels of task demand than will young adults. Thus, it is possible that in prior studies where age-invariant prefrontal monitoring effects were reported (e.g., de Chastelaine et al, 2016; Wang et al, 2015), task demands were such that monitoring did not run up against a resource limit in either young or older participants.

In the present study, we examined this proposal by using TMS to temporarily disrupt the right DLPFC, with the aim of reducing the neurocognitive resources available to support post-retrieval monitoring. Young and older adults studied visually presented word pairs followed by an associative recognition task. Offline TMS was administered to the right DLPFC or a control site (vertex) immediately prior to the retrieval phase. The right DLPFC was selected as the TMS target as the region has been previously reported to give rise to robust age-invariant retrieval monitoring effects in fMRI studies (de Chastelaine et al., 2016; Horne et al., 2020; Wang et al., 2015) and is readily accessible to administration of TMS.

Test items on the associative recognition test comprised pairs of ‘intact’ (studied together), ‘rearranged’ (studied on different trials), and ‘new’ (unstudied) word pairs. Word pairs endorsed as rearranged are thought to impose heavier demands on retrieval monitoring than word pairs endorsed as intact, because of the need to resolve the conflict between the familiarity of the individual test items and the novelty of the item association (de Chastelaine et al., 2016; Horne et al., 2020; see also Achim & Lepage, 2005). We predicted that TMS administered to the right DLPFC would lead to reduced associative recognition accuracy, and that this effect would be most prominent in older adults by virtue of their being closer to the ‘crunch point’ for monitoring than younger adults (see above). We further predicted that the effect would be driven largely by an increase in associative false alarms (rearranged pairs incorrectly endorsed as intact), under the assumption that rearranged items impose a heavier monitoring demand and hence are more vulnerable to the consequences of a monitoring failure (cf. Schacter et al., 1996). In addition, we anticipated that TMS administered to the right DLPFC might affect the response times (RT) associated with associative false alarms, and that this effect would be more pronounced in older adults. We were however agnostic as to the direction of any potential RT effects. One possibility is that the depletion of resources available to support retrieval monitoring will lead to a truncation of monitoring, and hence to faster RTs than those in the control condition (cf. Horne et al, 2020). Alternately, participants might attempt to compensate for the relative lack of resources by prolonging the engagement of monitoring processes, leading to a relative slowing of RTs.

## 2. Methods

### 2.1. Participants

Twenty-one healthy young adults and 18 healthy older adults volunteered and were monetarily compensated $30/hour for their time. All participants were right-handed, had normal or corrected-to-normal vision, and spoke fluent English before the age of five. Each participant had a structural MRI scan available from participation in a prior fMRI study. Self-reported TMS exclusion criteria followed safety guidelines (Rossi et al., 2009) and included potential pregnancy, prior or current neurological illness, current use of psychoactive medications, prior head injury or concussion, frequent or severe headaches/migraines, a personal or family history of seizure or epilepsy, and current use of prescription or over-the-counter medications associated with an increased risk of seizure. Additionally, prior to undergoing TMS, all participants acknowledged that they had not consumed alcohol in the previous 24 hours.

One young adult and two older adults voluntarily withdrew from the study prior to completing all cycles of the TMS protocol and their data were dropped from subsequent analyses. This resulted in a final sample of 20 healthy young adults (mean age: 23.5 years, range: 19-30 years, 12 female) and 16 healthy older adults (mean age: 69.6 years, range: 65-75 years, 6 female). A post-hoc power analysis conducted using G*Power (Faul et al., 2007) indicated that this sample size was sufficient to detect an effect size of f = .15 for the predicted age x TMS target interactions on recognition accuracy (β = .83) and false alarm rate (β = .87).

### 2.2. Neuropsychological Testing

All participants completed a neuropsychological test battery on a separate day consisting of the Mini-Mental State Examination (MMSE), the California Verbal Learning Test-II (CVLT; Delis et al., 2000), Wechsler Logical Memory Tests 1 and 2 (Wechsler, 2009), Trail Making tests A and B (Reitan & Wolfson, 1985), the Symbol Digit Modalities test (SDMT; Smith, 1982), the F-A-S subtest of the Neurosensory Center Comprehensive Evaluation for Aphasia (Spreen & Benton, 1977), the Wechsler Adult Intelligence Scale–Revised subtests of forward and backward digit span (Wechsler, 1981), category fluency (Benton, 1968), and Raven’s Progressive Matrices (List 1, Raven et al., 2000). In addition, participants completed the Wechsler Test of Adult Reading (WTAR, Wechsler, 2001) or its revised version, the Wechsler Test of Premorbid Functioning (TOPF; Wechsler, 2011).

Participants were excluded from entry into the study if they scored < 27 on the MMSE, < 1.5 SDs below age-appropriate norms on any memory test, < 1.5 SDs below age norms on any two other tests, or if their estimated full-scale IQ was less than 100. These criteria were employed to minimize the likelihood of including individuals with mild cognitive impairment. Results from the neuropsychological test battery are presented in Table 1. Note that the pattern of scores is typical of that reported for cross-sectional studies of cognitively healthy young and older adults (e.g., de Chastelaine et al., 2016; Wang et al., 2015), with higher scores for the young adults on tests involving episodic memory, processing speed, and fluid intelligence.

**Table 1.** Demographic and mean (with SD) neuropsychological test scores.

|  | Younger Adults | Older Adults | *p*-value |
| --- | --- | --- | --- |
| N | 20 | 16 |  |
| Sex (M/F) | 8/12 | 10/6 |  |
| Age | 23.11 (4.01) | 69.63 (3.07) |  |
| Years of education | 25.26 (1.91) | 16.88 (2.80) | .062 |
| MMSE | 28.40 (3.45) | 28.94 (1.00) | .515 |
| CVLT Short Delay - Free | 13.80 (4.38) | 10.94 (3.04) | .027* |
| CVLT Short Delay - Cued | 12.80 (3.04) | 12.56 (2.53) | .800 |
| CVLT Long Delay - Free | 12.98 (3.32) | 11.38 (2.87) | .131 |
| CVLT Long Delay - Cued | 13.08 (3.20) | 12.31 (2.44) | .424 |
| CVLT Recognition - Hits | 14.98 (3.22) | 15.25 (.93) | .720 |
| CVLT Recognition - False Alarms | 0.73 (1.41) | 2.50 (2.63) | .024* |
| Logical Memory I | 32.00 (8.16) | 30.06 (5.58) | .405 |
| Logical Memory II | 28.50 (6.15) | 27.19 (5.36) | .499 |
| SDMT | 60.75 (15.43) | 50.5 (8.08) | .016* |
| Digit Span | 19.30 (4.12) | 18.50 (2.88) | .498 |
| Trails A | 20.53 (8.08) | 26.73 (9.27) | .040* |
| Trails B | 44.45 (13.03) | 63.55 (22.98) | .007* |
| F-A-S | 46.43 (17.82) | 48.06 (12.61) | .749 |
| Category Fluency (Animals) | 24.65 (6.48) | 22.25 (4.43) | .197 |
| WTAR/TOPF<sup>1</sup> | 114.94 (5.40) | 114.81 (8.67) | .959 |
| Raven's (List 1) | 10.95 (.91) | 8.94 (1.88) | < .001* |

*Note:* Significant differences between age groups are denoted by an asterisk. <sup>1</sup>Presented as a standardized score computed from WTAR and TOPF raw scores. MMSE = Mini-Mental State Examination; CVLT = California Verbal Learning Test; SDMT = Symbol Digit Modalities Test; WTAR = Wechsler Test for Adult Reading; TOPF = Wechsler Test of Premorbid Functioning.

### 2.3. MRI Acquisition and Preprocessing

MRI data were acquired with a 3T Philips Achieva MRI scanner (Philips Medical Systems, Andover, MA, USA) equipped with a 32-channel receiver head coil. T1-weighted structural images (MPRAGE sequence, 240 × 240 matrix, 1mm isotropic voxels) were acquired. Each T1 image was normalized to an independent sample-specific template derived from 36 young and 64 older adult T1 images using Statistical Parametric Mapping (SPM12) software. The transformation matrix was then used to back-project a DLPFC ROI into each participant’s native space using neuronavigation software (see TMS procedure).

### 2.4. Associative Memory Task

The experiment proper consisted of two study-test cycles corresponding to the right DLPFC and vertex TMS stimulation conditions, the order of which was counterbalanced across participants. A 45 minute stimulation ‘washout’ period separated the two study-test cycles to allow any residual stimulation effects to dissipate (Huang et al., 2005).

#### 2.4.1. Stimuli

Cogent 2000 software (www.vislab.ucl.ac.uk/cogent.php) implemented in Matlab 2012b (www.mathworks.com) was used for stimulus presentation and response collection. The stimuli were presented on a 30” Dell LCD monitor with a screen resolution of 1024 × 768 pixels at a viewing distance of ~100 cm. All critical stimuli for the study and test phases were presented in white Helvetica 30-point font on a black background. Study and test items subtended a vertical visual angle of 2.7° and a maximum horizontal visual angle of 6.2°.

Experimental stimuli comprised 240 semantically unrelated word pairs. All words were concrete nouns between three and nine letters in length, with a mean frequency per million of 8.28 (SD = 31.69, range 0.04 to 479.92) obtained from the SUBTLEX-US corpus (Brysbaert & New, 2009). Concreteness ratings ranged between 3 and 5 (*M* = 4.67, *SD* = .29) on a scale from 1 (most abstract) to 5 (most concrete) (Brysbaert et al., 2014). Separate stimulus lists were created for yoked young and older adult pairs. Two sets of 90 randomly selected word pairs served as the critical stimuli for the study phase. An additional two sets of 120 word pairs served as the critical stimuli for the test phase. Each test list included 60 ‘intact’ pairs (words presented together at study), 30 ‘rearranged’ pairs (words presented on different trials at study), and 30 ‘new’ pairs (words not encountered at study). All test pairs were inspected by the second author to ensure they were not semantically related. Study and tests lists were assigned to one of the two TMS stimulation conditions (right DLPFC, vertex). An additional 24 word pairs were used as practice stimuli. The practice study phase comprised 18 word pairs, and the practice test phase comprised 24 pairs (12 ‘intact’ pairs, 6 ‘rearranged’ pairs, and 6 ‘new’ pairs).

#### 2.4.2. Study Phase

Word pairs were presented for a duration of 2000 ms and were preceded by a red fixation cross for 500 ms. The presentation of each word pair was followed by a white fixation cross for 1000 ms, giving a response window of 3000 ms per trial. The study phase for each TMS stimulation condition (DLPFC, vertex) was administered as a single block and lasted approximately 6 minutes (see Figure 1 for task schematic). Encoding was intentional, as participants were aware of the subsequent associative recognition test and trained on both study and test phases before beginning the experiment. Study words were presented simultaneously, one above and one below fixation. The task was to judge which of the two objects denoted by the words would more likely ‘fit’ into the other and to respond via a button press. To encourage relational encoding of the word pairs, participants were instructed to focus on imagining a scenario (constructing a vivid visual image or verbal story) to determine which item would fit into the other. Participants were instructed to respond as quickly as possible without sacrificing accuracy.

**Figure 1.**
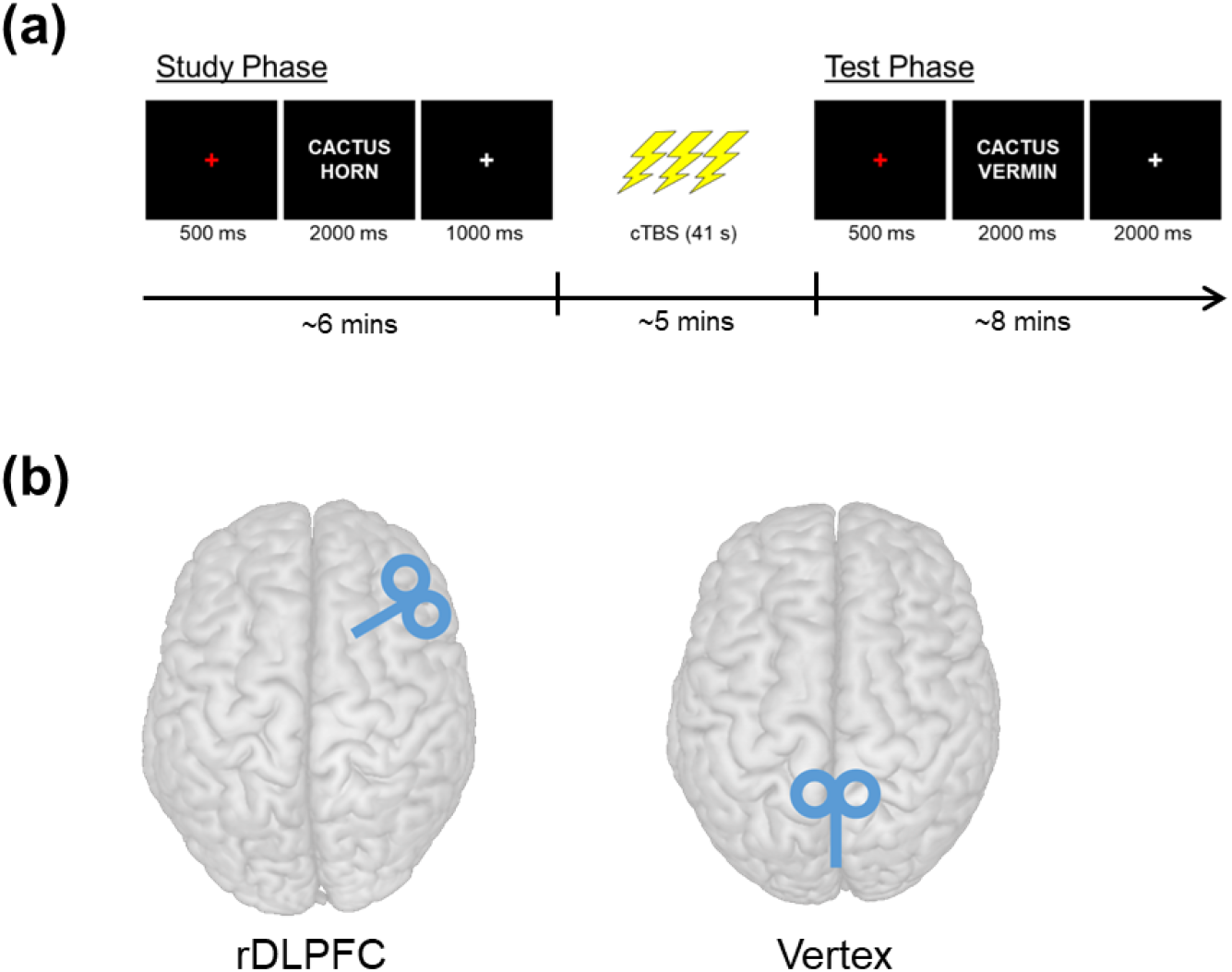
(a) Timing and schematic of the associative memory task. (b) TMS targets in the right dorsolateral prefrontal cortex (left) and vertex (right).

**Figure 2.**
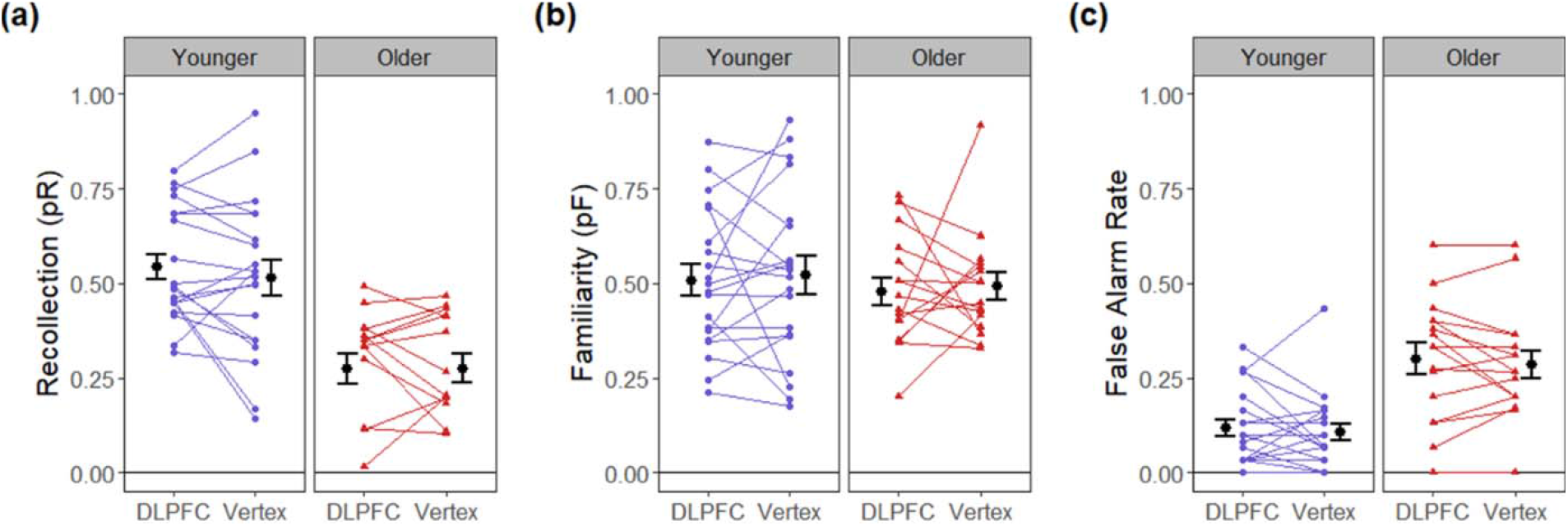
Estimates of (a) recollection, (b) familiarity, and (c) false alarm rates for the DLPFC and vertex targets in young and older adults. Error bars reflect the standard error of the mean.

**Figure 3.**
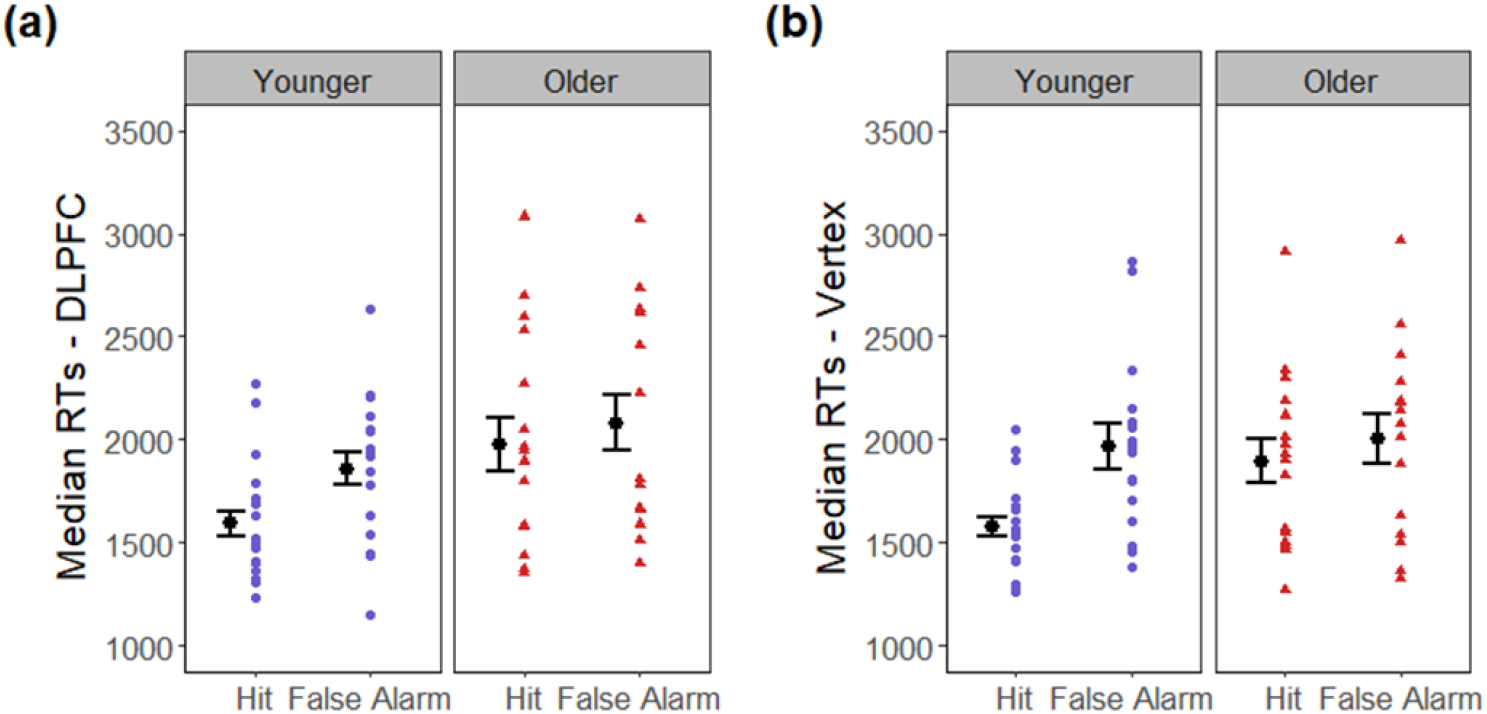
Median test phase response times (RTs) for associative hits and associative false alarms for young and older adults in (a) right DLPFC and (b) vertex TMS target conditions. Error bars reflect the standard error of the mean.

#### 2.4.3. Test Phase

The test phase began approximately 5-10 minutes after completion of the study phase, immediately following the TMS procedure. Word pairs during the test phase were presented for a duration of 2000 ms and were preceded by a red fixation cross for 500 ms. The presentation of each word pair was followed by a white fixation cross for 2000 ms, giving a response window of 4000 ms per trial. The test phase for each TMS stimulation condition (DLPFC, vertex) was administered as a single block and lasted approximately nine minutes (Figure 1A). Test words were presented simultaneously, one above and one below fixation.

For the associative memory task, participants were required to judge whether each word pair was ‘intact’, ‘rearranged’, or ‘new’. Participants were instructed to respond ‘intact’ when they could recall with high confidence and specificity that the two items had been studied together, and to respond ‘rearranged’ when the words were recognized as having been studied but there was either no memory of their having been studied together, or the memory was uncertain. Participants were informed that the test list did not include mixed pairs of new and studied words. They were however instructed to respond ‘new’ whenever they could recognize only one of the words as studied, or when both words were judged to be unstudied. Participants again indicated their responses via a button press; they were instructed to respond as quickly as possible without sacrificing accuracy.

### 2.5. TMS Procedure

#### 2.5.1. Equipment and co-registration

TMS was delivered using a Magstim Super Rapid2 stimulator equipped with a 70-mm figure-of-eight coil. Frameless stereotaxy was used to co-register participants to their MRI scans and to align the coil over the target stimulation sites using Brainsight (Rogue Research; www.rogue-research.com/) software equipped with a Polaris infrared camera. During co-registration and TMS, each participant’s head was stabilized with a padded headrest to minimize movement.

Four anatomical landmarks were identified on the skin reconstruction of a participant’s MRI scan: nasion, nose tip, and bilateral pre-auricular points. The nose tip was excluded if it was outside of the field-of-view of the structural MRI, leaving only three points for co-registration. Co-registration was performed by identifying the relevant anatomical landmarks on the participant’s head using a pointer; this allowed us to track the location of the coil in real-time relative to the ROIs targeted for stimulation for each participant. Reflective markers attached to the TMS coil and each participant’s head (via an elastic velcro strap) were used to monitor coil position and head location, respectively, using the infrared camera.

#### 2.5.2. Motor Threshold

For each participant, a motor threshold was determined using single pulse TMS over the left motor cortex (visually identified on the structural MRI) using a procedure identical to that described in detail by Koen and colleagues (2018). In brief, a motor threshold was defined as the lowest stimulator intensity that produced visible muscle contractions in the right hand in 5 out of 10 pulses. In a minority of participants, motor threshold was estimated with motor evoked potentials in lieu of visible contractions. The mean motor threshold was 62% (SD = 8%) of maximum stimulator output in younger adults and 65% (SD = 5%) in older adults. Stimulator intensity for continuous theta-burst stimulation (cTBS, Huang et al., 2005; see below) was set nominally at 80% of motor threshold with the constraint that it never exceeded 45% of maximum stimulator output. This constraint was due to a hardware limitation of the Rapid2 stimulator. Thus, cTBS threshold was set at 45% for participants in whom no motor response was identified, or if 80% of the identified motor thresholds was above the 45% cutoff. This resulted in a cTBS threshold that was set at the 45% maximum threshold in 17 of 20 young adults and all older adults. The resultant stimulation intensity in young (*M* = 44%, *SD* = 2%) and older (*M* = 45%, *SD* = 0%) adults did not significantly differ between the two age groups (*t*_(19)_ = 1.75, *p* = .096). These intensities corresponded to stimulation that averaged 72% (SD = 7%, range 58-81%) of motor threshold in the young sample, and 69% (SD = 5%, range 61-75%) in the older group. These means also did not differ between the groups (*t*_(21.96)_ = −1.17, *p* = .253).

#### 2.5.3. Stimulation Protocol

Offline cTBS was delivered to either the right DLPFC or vertex target immediately following each study phase and hence prior to the beginning of the succeeding test phase. The cTBS protocol involved delivering 50 Hz trains of three TMS pulses every 200 ms for 40 s (600 total pulses) at 80% resting motor threshold (see above). Similar cTBS protocols have been employed to suppress cortical excitability in prior studies of episodic memory (Berkers et al., 2017; Bonnici et al., 2018; Marin et al., 2018; Ryals et al., 2016; Yazar et al., 2014, 2017). The order of the DLPFC and vertex targets was counterbalanced across subjects. The right DLPFC target (MNI coordinates: × = 48, y = 32, z = 28) was the same as that defined in a previously published experiment using the associative memory design (de Chastelaine et al., 2016). The vertex was selected for the control stimulation condition (Koen et al., 2019; Sandrini et al., 2003; Thakral et al., 2017; Yazar et al., 2014, 2017), and was identified in each participant as the intersection between the central sulcus and longitudinal fissure.

### 2.6. Statistical Analyses

Statistical analyses were carried out using R software (R Core Team, 2017). ANOVAs were conducted using the *afex* package (Singmann et al., 2016) and the Greenhouse-Geisser procedure (Greenhouse & Geisser, 1959) was used to correct degrees of freedom for non-sphericity when necessary. Post-hoc tests on significant effects from the ANOVAs were conducted using the *emmeans* package (Lenth, 2018). Results were considered significant at *p* < .05. We also report results from Bayesian repeated-measures ANOVAs using JASP software (version 0.13.1.0). Specifically, we computed Bayes Factors (BF) corresponding to evidence in favor of the effects of interest (e.g., the age × TMS target interaction contrasts) relative to the null model. We report these values as BF_10_, with values < 1 indicating relatively greater evidence favoring the null hypothesis. The BF_10_ for ANOVA main effects and interactions were computed using the ‘BF Inclusion’ (reported here as BF_10_ for consistency) values output by JASP with the ‘Across Matched Models’ option.

## 3. Results

A summary of associative memory performance and median RTs for each age group and TMS target is shown in Table 2. The proportion of ‘intact’, ‘rearranged’, and ‘new’ test responses for each of the different pair types is reported in Table 3. The critical trials used in the following analyses included associative hits (intact pairs correctly endorsed as intact), associative misses (intact pairs incorrectly endorsed as rearranged), and associative false alarms (rearranged pairs incorrectly endorsed as intact). Estimates of recollection accuracy (pR) were indexed as the difference between the proportion of associative hits and associative false alarms (de Chastelaine et al., 2016; Horne et al., 2020) using the formula:

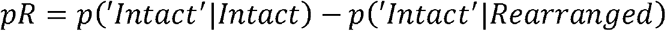

**Table 2.** Mean (with SD) of associative memory performance and median RT as a function of age group and TMS target

| Measure | Younger |  | Older |  |
| --- | --- | --- | --- | --- |
|  | DLPFC | Vertex | DLPFC | Vertex |
| Recollection (pR) | .54 (.15) | .51 (.21) | .28 (.16) | .28 (.15) |
| Familiarity (pF) | .51 (.19) | .52 (.23) | .48 (.15) | .49 (.14) |
| False Alarm Rate | .12 (.10) | .11 (.10) | .30 (.16) | .29 (.15) |
| *Median RTs* |  |  |  |  |
| Associative Hits | 1613 (301) | 1584 (231) | 1927 (512) | 1919 (427) |
| Associative False Alarms | 1825 (295) | 1967 (435) | 2078 (522) | 2004 (469) |

**Table 3.** Mean proportion (with SD) of responses to intact, rearranged, and new test pairs as a function of age group and TMS target.

| Trial Type | Response | Younger |  | Older |  |
| --- | --- | --- | --- | --- | --- |
|  |  | DLPFC | Vertex | DLPFC | Vertex |
| Intact | Intact | .66 (.14) | .62 (.19) | .58 (.20) | .56 (.19) |
|  | Rearranged | .21 (.10) | .22 (.11) | .30 (.17) | .30 (.16) |
|  | New | .13 (.09) | .16 (.12) | .13 (.06) | .14 (.08) |
| Rearranged | Intact | .12 (.10) | .11 (.10) | .30 (.16) | .29 (.15) |
|  | Rearranged | .69 (.16) | .63 (.18) | .49 (.14) | .49 (.18) |
|  | New | .19 (.14) | .26 (.14) | .21 (.11) | .23 (.13) |
| New | Intact | .01 (.02) | .01 (.03) | .03 (.05) | .04 (.04) |
|  | Rearranged | .12 (.08) | .11 (.11) | .18 (.13) | .17 (.13) |
|  | New | .87 (.09) | .88 (.12) | .79 (.15) | .80 (.13) |

Estimates of familiarity strength (pF) were estimated under the independence assumption (Yonelinas & Jacoby, 1995) using the formula:

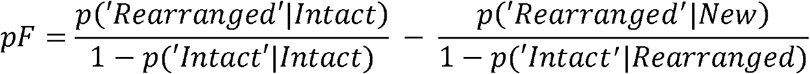

Estimates of recollection (pR) and familiarity (pF) were entered into separate 2 (age group: younger, older) × 2 (TMS target: DLPFC, vertex) ANOVAs. There was a significant main effect of age on pR (*F*_(1,34)_ = 23.36, *p* < 10^−4^, partial-η^2^ = .41, BF_10_ = 552.83) which was driven by reduced recollection accuracy in older adults relative to their younger counterparts. The main effect of TMS target on pR was not significant (*F*_(1,34)_ = 0.47, *p* = .498, partial-η^2^ = .01, BF_10_ = .31), nor was there a significant age × TMS target interaction (*F*_(1,34)_ = 0.50, *p* =.484, partial-η^2^ = .01, BF_10_ = .38). The analysis of pF failed to reject the null hypothesis for the main effects of age (*F*_(1,34)_ = 0.31, *p* = .579, partial-η^2^ = .009, BF_10_ = .40) and TMS target (*F*_(1,34)_ = 0.17, *p* = .679, partial-η^2^ = .005, BF_10_ = .27), or the age × TMS target interaction (*F*_(1,34)_ = 0.02, *p* =.902, partial-η^2^ = .001, BF_10_ = .29).

An ANOVA performed on the proportions of associative false alarms (see Introduction) revealed a significant main effect of age (*F*_(1,34)_ = 20.54, *p* < 10^−4^, partial-η^2^ = .38, BF_10_ = 236.01) which was driven by an elevated false alarm rate in older relative to young adults. The main effect of TMS target was not significant (*F*_(1,34)_ = 0.80, *p* = .378, partial-η^2^ = .02, BF_10_ = .34), nor was there a significant age × TMS target interaction (*F*_(1,34)_ = 0.02, *p* =.902, partial-η^2^ = .001, BF_10_ = .33).

To investigate the effect of TMS condition on retrieval response times, we entered the median test phase RTs for associative hits and associative false alarms into a 2 (age) × 2 (TMS target) × 2 (memory: hit, false alarm) ANOVA. This analysis revealed a significant three-way interaction (*F*_(1,29)_ = 4.77, *p* = .037, partial-η^2^ = .14, BF_10_ = 0.922). To unpack these results, we performed subsidiary 2 (age) × 2 (memory) ANOVAs on test phase RTs for the right DLPFC and vertex TMS conditions, respectively. The analysis of RTs in the right DLPFC condition revealed a significant main effect of age (*F*_(1,32)_ = 4.32, *p* = .046, partial-η^2^ = .12) which was driven by faster RTs for young compared to older adults, as well as a main effect of memory response (*F*_(1,32)_ = 17.89, *p* = .0002, partial-η^2^ = .36) which was driven by faster RTs for associative hits compared to associative false alarms. The age × response interaction was not significant (*p* = .233, partial-η = .04). An analogous analysis of RTs in the vertex condition revealed a significant age × memory interaction (*F*_(1,29)_ = 11.88, *p* = .002, partial-η^2^ = .29) which was driven by significantly faster RTs for hits compared to false alarms in young adults (*t*_(29)_ = 6.35, *p* <.0001, BF_10_ = 26.66) but not older adults (*t*_()_ = 1.36, *p* =.186, BF_10_ = .40).

## Discussion

In the present study, we investigated whether healthy older adults were more vulnerable than younger adults to an experimental manipulation that was intended to deplete the availability of neurocognitive resources associated with post-retrieval monitoring. To this aim, we delivered offline TMS to the right DLPFC and a control site (vertex) immediately prior to completion of an associative recognition test. Collapsed across the two TMS target conditions, recognition accuracy was significantly lower in older relative to young adults. By contrast, estimates of familiarity strength did not reliably differ between the two age groups. This pattern of results is consistent with numerous prior findings of attenuated recollection accompanied by relatively preserved familiarity in healthy older adults (e.g., de Chastelaine et al., 2016; de Chastelaine et al., 2017; Wang et al., 2015; for review, see Koen & Yonelinas, 2014; Schoemaker et al., 2014).

We predicted that TMS administered to the right DLPFC would lead to reduced associative recognition accuracy, and that this effect would be more prominent in older adults. We also predicted that the effect would be driven largely by an increase in associative false alarms. Our results however provide no evidence that TMS administered to the right DLPFC had any impact on associative recognition accuracy in either age group. These null results were supported by Bayes Factors indicating moderate to strong evidence in favor of a null effect of TMS target condition on associative memory performance (the null model was ~2.5 times more likely than the alternative model). The results therefore suggest that any depletion of cognitive resources dependent on the right DLPFC resulting from TMS was not sufficient to impact the accuracy of associative recognition judgments regardless of age.

As was noted in the Introduction, we selected the right DLPFC as a TMS target because the region has been previously reported to give rise to robust age-invariant retrieval monitoring effects (de Chastelaine et al., 2016; Horne et al., 2020; Wang et al., 2015) and is readily accessible to administration of TMS. However, monitoring effects have also been consistently reported in other regions, most notably in the anterior cingulate cortex and left DLPFC (de Chastelaine et al., 2016; Horne et al., 2020; Wang et al., 2015). These findings raise the possibility that redundancy across the different regions allowed for sufficient neurocognitive resources to compensate for depletion arising from TMS of the right DLPFC.

We also predicted that TMS administered to the right DLPFC would affect RTs for associative false alarms, and that this effect would be most prominent in older adults. This prediction too was unfulfilled. However, we did identify a significant effect of TMS on RTs to test items, albeit in young rather than older participants. Whereas RTs were robustly shorter for associative hits than false alarms in the vertex stimulation condition in the young group, this difference was absent after TMS to the right DLPFC. By contrast, older adults failed to demonstrate an RT advantage for hits in either stimulation condition. Thus, TMS administered to the right DLPFC appears to have shifted the patterning of the RTs for hits and false alarms in the young adults so that it resembled the pattern evident in the older group. We conjecture that this effect might reflect an impact of TMS on the confidence of the young adults’ ‘intact’ judgments to intact and rearranged pairs. By this argument, while confidence was higher in these participants (but not in older adults) for correct than for incorrect judgments in the control condition (reflected in the differential RTs), this was not the case after TMS to the DLPFC. We stress however that these results were unanticipated, and that they most decidedly merit replication. That being said, while not supporting our initial predictions, the results do provide evidence that the TMS intervention impacted some aspects of task-related behavior. Thus, they provide reassurance that the null results of TMS on associative recognition accuracy did not stem from a blanket failure of the TMS protocol to modulate neural function.

The present null findings converge with those from a study employing a quite different strategy to examine the age-related ‘resource limitation hypothesis’ motivating the present study. Horne et al. (2020) employed a dual-task procedure in which an associative recognition test was paired with a secondary tone detection task in an effort to limit the neurocognitive resources available to support post-retrieval monitoring. While the manipulation disproportionately impaired the associative recognition performance of older relative to young participants, fMRI correlates of retrieval monitoring, including those in right DLPFC, were unaffected in both age groups. Thus, as in the present case, these fMRI findings suggest that the monitoring operations supported by this region are relatively resistant to a manipulation aimed at disrupting these operations by limiting access to domain-general processing resources.

A potential limitation of the present study concerns the spatial extent of TMS. Right DLPFC is a sizeable and, it has been argued (Duncan, 2001), near-equipotential cortical region, and the relatively focal stimulation provided by TMS (~15 cm^2^, Deng et al., 2013) might have failed to disrupt a large enough proportion of the tissue capable of supporting monitoring. Future studies using methods that have the potential to disrupt a broader expanse of the DLPFC, such as transcranial direct-current stimulation, might yield different results (e.g., Gaynor & Chua, 2019). Another potential limitation stems from our use of a group-level fMRI monitoring effect (de Chastelaine et al., 2016) to identify right DLPFC TMS targets at the single participant level. It is possible that our null findings arose due to inter-subject variability in the locus of the critical DLPFC region necessary for effective monitoring operations. We note however that prior studies have reported successful TMS interventions using a similar approach to targeting TMS (Thakral et al., 2017; Yazar et al., 2014; 2017). A third limitation is that, although we aimed for a stimulation intensity of 80% of motor threshold, the majority of our participants were stimulated at a lower intensity due to a hardware limitation (see methods). Thus, our null results might reflect the employment of an inadequate stimulation intensity. While we cannot rule out this possibility it is noteworthy that Bonnici et al. (2018) and Yazar et al. (2014; 2017) employed stimulation intensities of a similar magnitude to those employed here (70% motor threshold) to study the role of the angular gyrus in memory retrieval and reported positive findings. Moreover, it has been reported that the relationship between the intensity of cTBS stimulation and degree cortical suppression is complex and possibly non-linear, at least in some participants, casting doubt on the assumption that the behavioral effects of such stimulation should correlate positively with stimulation intensity (Sasaki et al., 2018). Further work is needed to establish the stimulation parameters that have the greatest likelihood of impacting episodic memory processes (Yeh & Rose, 2019; Widhalm & Rose, 2019).

In conclusion, we found that TMS to the right DLPFC had no detectable impact on associative memory performance in young or older adults. These results suggest that stimulation of the right DLPFC was insufficient to deplete the neurocognitive resources available to support post-retrieval monitoring.

